# Direct medical and non-medical costs of a one-year care pathway for early breast cancer: results of a French multicenter prospective study

**DOI:** 10.1101/514182

**Authors:** Delphine Héquet, Cyrille Huchon, Anne-Laure Soilly, Bernard Asselain, Hélène Berseneff, Caroline Trichot, Alix Combes, Karine Alves, Thuy Nguyen, Roman Rouzier, Sandrine Baffert

**Affiliations:** Department of Surgical Oncology, Institut Curie-René Huguenin, 35 rue Dailly, 92210, St Cloud, France.; Department of Gynecology, Poissy-St Germain hospital, 10 Rue du Champ Gaillard, 78300, Poissy, France.; Health Economics Department, Bourgogne Franche-Comté University, EA 7467 Dijon, France/ CHU Dijon Bourgogne, Délégation à la Recherche Clinique et à l’Innovation, USMR, F-21000 Dijon; Department of Biostatistics, Institut Curie, 26 rue d’Ulm, 75005, Paris, France.; Department of Gynecology, René Dubos Hospital, 6, avenue de L’Ile de France, 95303, Pontoise, France.; Department of Gynecology, Antoine Béclère Hospital, 157 rue de la Porte de Trivaux, 92140, Clamart, France.; Department of Gynecology, André Mignot Hospital, 50 rue Berthier, 78000, Versailles, France.; Department of Gynecology, Argenteuil Hospital, 69 Rue Lt Colonel Prudhon, 95100, Argenteuil, France.; Department of Gynecology, Louis Mourier Hospital, 178 rue des Renouillers, 92700, Colombes, France.; Health Economics Department, Institut Curie, 26 rue d’Ulm, 75005, Paris, France/CEMKA-EVAL, 43 Boulevard du Maréchal Joffre, 92340 Bourg-La-Reine, France.

## Abstract

**Introduction:** The organization of health care for breast (BC) constitutes a public health challenge to ensure quality of care, while also controlling expenditure. Few studies have assessed the global care pathway of early BC patients, including a description of direct medical costs and their determinants.

**Methods:** OPTISOINS01 was a multicenter, prospective, observational study including early BC patients from diagnosis to one-year follow-up. Direct medical costs (in-hospital and out-of-hospital costs, supportive care costs) and direct non-medical costs (transportation and sick leave costs) were calculated by using a cost-of-illness analysis based on a bottom-up approach. Resources consumed were recorded *in situ* for each patient, using a prospective direct observation method.

**Results:** Data from 604 patients were analyzed. Median direct medical costs of 1 year of management after diagnosis in operable BC patients were €12,250. Factors independently associated with higher direct medical costs were: diagnosis on the basis of clinical signs, invasive cancer, lymph node involvement and conventional hospitalization for surgery. Median sick leave costs were €8,841 per patient and per year. Chemotherapy was an independent determinant of sick leave costs (€3,687/patient/year without chemotherapy versus €10,706 with chemotherapy). Forty percent (n=242) of patients declared additional personal expenditure of €614/patient/year. No drivers of these costs were identified.

**Conclusion:** Initial stage of disease and the treatments administered were the main drivers of direct medical costs. Direct non-medical costs essentially consisted of sick leave costs, accounting for one-half of direct medical costs for working patients. Out-of-pocket expenditure had a limited impact on the household.

## INTRODUCTION

Breast cancer (BC) is the most common cancer in women in France with more than 54,000 new cases and 12,000 deaths per year [1]. BC therefore constitutes a public health challenge due to the need to organize health care while ensuring quality of care and controlling expenditure. Moreover, BC is a heterogeneous disease, in which treatments, clinical practices and costs can vary substantially. A care pathway is defined as a patient-focused global care that addresses temporal (effective and coordinated management throughout the illness) and spatial issues (treatment is provided near the health territory in or around the patient’s home) [2]. Studying care pathways therefore implies focusing on a specific health care territory and identifying health care services and resources consumed in the various health care structures. The BC care pathway is complex, involving several structures, disciplines, actors, technical, organizational and medical innovations. Analysis of BC global care pathways, including health care, support and treatment, is needed to help patients, heath care professionals and policy-makers. Clinical pathways can then be used as a tool to standardize processes and improve quality of care, as previously conducted in other fields of medicine: congestive heart failure, stroke, asthma, or deep vein thrombosis [3–6]. Most studies on BC care pathways have focused on a specific phase of treatment, such as surgery [7,8] or radiotherapy [9] and few studies have assessed the complete and global care pathway of early BC patients, including treatment, support and innovative organizations, such as the use of outpatient surgery.

The costs of BC are also underestimated. Most published health economics, cost-effectiveness or cost-utility studies have focused on a specific drug [10], surgical procedure [11] or examination [12,13]. Few studies have assessed the overall direct costs of one-year management, including medical and non-medical costs, in order to elucidate the distribution of costs during the various phases of health care (diagnosis, treatment, follow-up), which would be useful for policymakers to inform decisions on health care funding.

The aims of this multicenter prospective study were to describe care pathways of BC patients in a geographic territory and to calculate the global direct costs of early stage BC during the first year following diagnosis.

## MATERIALS AND METHODS

OPTISOINS01 was a multicenter, prospective, observational study conducted in patients from a determined regional health territory around Paris. This study was approved by the French National ethics committee (CCTIRS Authorization No. 14.602 and CNIL DR-2014-167) for the research conducted at all participating hospitals. This study was registered with ClinicalTrials.gov (Identifier: NCT02813317). The design of this study has been previously published [14].

### Setting and population

The study was conducted in three departments of the Ile-de-France region (Hauts-de-Seine, Yvelines, and Val d’Oise), which cover 35% of the region’s population (total population: 11.9 million). This region has a population of 2.17 million women, 61% of whom are older than 45 years, with an incidence and mortality of BC higher than national rates. This territory was chosen due to its heterogeneity in terms of health care services provided and its variable medical density and health care facilities, related to the disparate urbanization and population incomes throughout the territory. Eight non-profit hospitals participated in the study: 3 teaching hospitals (TH), 4 general hospitals (GH) and 1 comprehensive cancer center (CCC). Patients included in the study had histologically confirmed, previously untreated, operable breast cancer, and were living in one of the three departments of the region studied. Informed consent was obtained from all individual participants included in the study.

### Breast cancer care pathways

As BC is a heterogeneous disease, depending on the stage and characteristics of the cancer, the study population was divided into three groups of patients. These groups corresponded to patients with homogeneous BC pathways, based on clinical relevance and the treatments administered. Group 1 was composed of patients treated by conservative breast surgery, without axillary dissection or chemotherapy. This group was used to identify factors associated with outpatient surgery, as axillary dissection (group 2) and radical mastectomy (group 3) may constitute a contraindication to outpatient surgery for some teams. Patients of group 2 were also treated by conservative breast surgery, but associated with either axillary dissection or chemotherapy. Group 3 included all patients treated by radical breast surgery.

### Costing methodology

The overall cost of the one-year early breast cancer pathway was determined by cost-of-illness analysis. Resources consumed were recorded *in situ* for each patient, using a prospective direct observation method [15]. In addition, each patient filled in a diary to provide information about sociodemographic status, outpatient health care consumption, and modes of work and social reintegration. The time horizon was the period from breast cancer diagnosis until one year following diagnosis. The pathway costs were calculated for the overall pathway, including four phases: the diagnosis phase (first examination with diagnosis of BC until surgery), surgery and post-surgery phase (from surgery to initiation of adjuvant therapy), adjuvant treatment phase (from initiation of first adjuvant therapy until the end of last adjuvant therapy, excluding hormone therapy) and the one-year follow-up phase (from the end of adjuvant therapy until one year after diagnosis). Costs were calculated from societal perspectives, regardless of the funding actor and were limited to direct costs (medical and non-medical). Direct costs, including direct medical direct (in-hospital and out-of-hospital costs, supportive care costs) and direct non-medical costs (transportation and sick leave costs) were assessed by a bottom-up method. Costs are expressed in 2016 euros (including VAT).

### Statistical analysis

For each cost component, the mean, standard deviation (SD), minimum, maximum and median were calculated for quantitative variables and frequencies and percentages were calculated for qualitative variables. Univariate and multivariate analysis was performed to establish relationships between patient characteristics, patient care and costs. Fisher’s exact test, Student’s t-test, Wilcoxon test or Mann-Whitney test were used to analyze these factors. Multivariate analysis was performed using a logistic regression model. Differences were considered significant for p<0.05. All statistical analyses were performed with R software [16].

## RESULTS

### Pathways characteristics and resources consumed

More than 750 patients were screened for participation in the study between January 2014 and January 2016 and 617 of them were included in the study. After monitoring and lost to follow-up, data for 604 patients were analysed and 303, 164 and 137 patients were assigned to group 1, group 2, and group 3, respectively (Figure 1).

**Figure 1:**
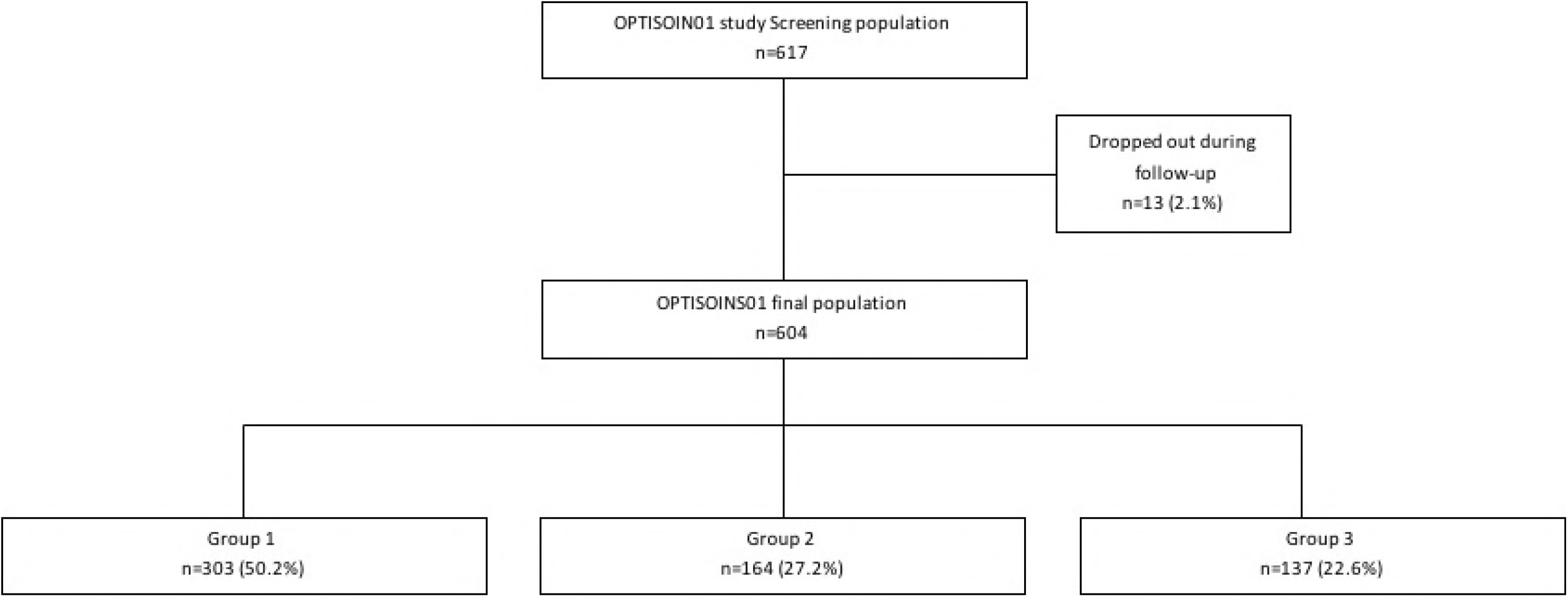
Flow chart of Optisoins01 study population

Median age was 58 years [27–94]. Most patients presented invasive BC (n=543, 90%) and 23% (n=142) had axillary lymph node involvement. Less than 40% of patients received adjuvant chemotherapy (n=235, 39%). Nearly one-half of patients were working at the time of diagnosis (n=297, 49%). Patient characteristics and treatments administered are presented in Table 1.

**Table 1:**
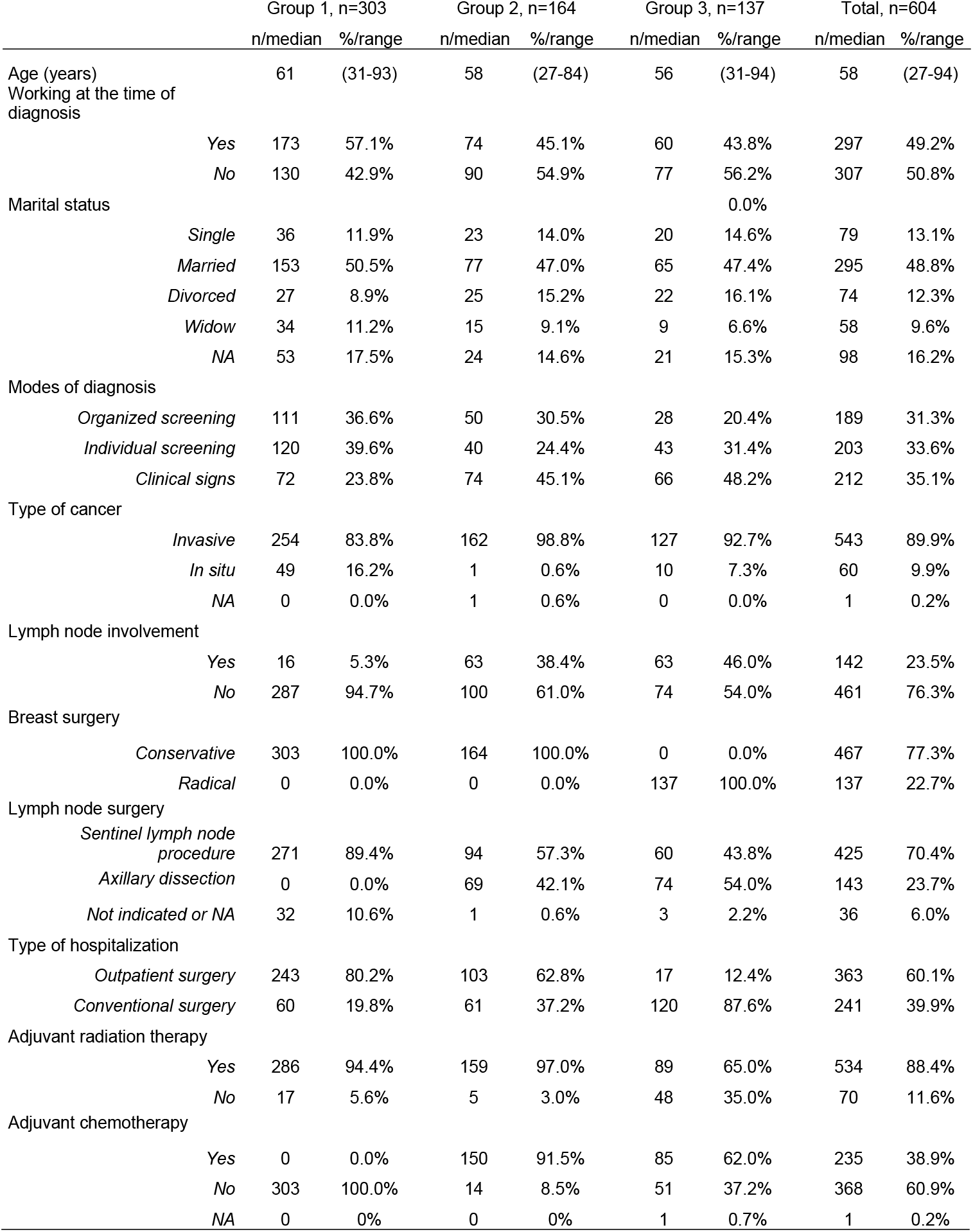
Patients, cancer and treatments characteristics, n=604

The majority of patients (n=447, 74%) were managed in a CCC, 14% in a GH (n=83) and 12% in a TH (n=12%). Median distance between residence and center was 12 km. Patients travelled longer distances to attend a CCC (18 km) than a GH or TH (6 km).

Among the 379 patients between the ages of 50 and 74 years (target population of systematic screening in France), 290 (77%) were diagnosed following a screening examination and 89 (23%) were diagnosed on the basis of clinical signs.

More than 80% of group 1 patients were managed by outpatient surgery (n=243, 80.2%). In the multiple linear regression model, 2 factors were associated with outpatient surgery in this group: being operated in a CCC (OR: 7.3, 95%CI[3.5-15.6], p<0.0001) and living in a couple (OR:2.6, 95%CI[1.3-5.4], p=0.007).

Resources consumed included medical consultations, imaging examinations for diagnosis (mammograms, breast ultrasound, breast Magnetic Resonance Imaging), biopsies, staging examinations (positron emission tomography-scanner, thoracoabdominal CT-scan, bone scan, chest radiograph, blood test), pre-treatment examinations, supportive care (psychology, nutrition, physiologist), surgery, hospitalizations, radiation therapy sessions, chemotherapy courses; for each phase of diagnosis (Table 2).

**Table 2:**
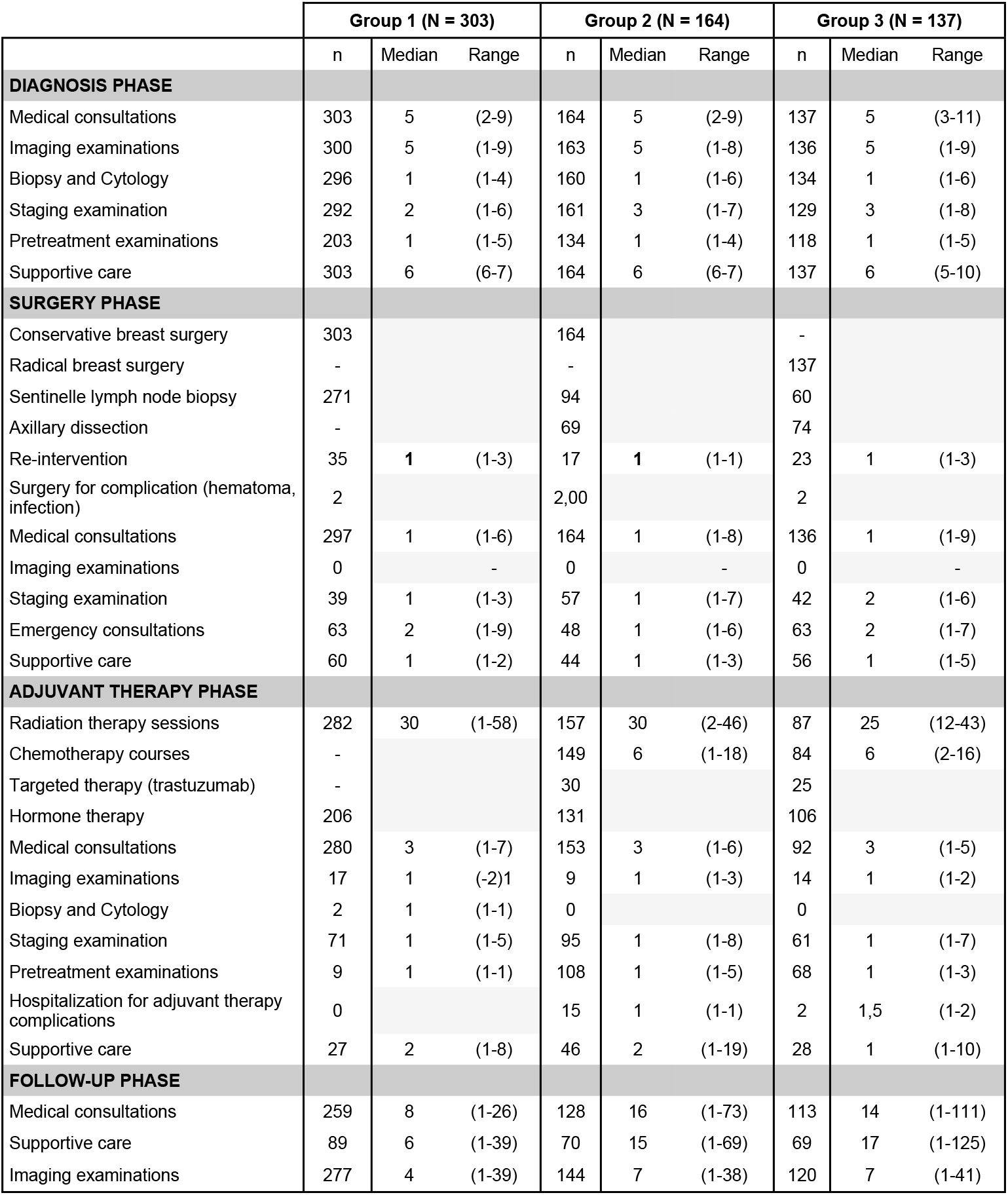
consumed resources (median and range are given for consuming patients only (n))

### Pathway cost

#### Direct medical costs from a National Health Insurance perspective

Median direct medical costs of 1 year of management from diagnosis of operable BC were €12,250 (€3,643-€55,340). Table 3 shows direct medical costs by phase and by group. Except for the follow-up phase, a statistically significant difference of these costs was observed between the 3 clinical pathway groups, with costs about 30% lower for group 1 compared to the other groups.

**Table 3:** Direct medical costs by phase of management and clinical pathway group (€)

|  | Group 1, n=303 |  | Group 2, n=164 |  | Group 3, n=137 |  | p |
| --- | --- | --- | --- | --- | --- | --- | --- |
|  | Median | Range | Median | Range | Median | Range |  |
| **Diagnosis** | **731** | (304-3,253) | **839** | (325-2,618) | **1,035** | (228-8,423) | <.0001 |
| **Surgery** | **3,427** | (2,445-17,808) | **3,530** | (2,445-11,560) | **5,993** | (3,590-14,959) | <.0001 |
| **Adjuvant therapy** | **6,538** | (0-11,773) | **9,982** | (111-35,428) | **8,004** | (0-32,297) | <.0001 |
| **Follow-up** | **666** | (0-3,155) | **666** | (0-2,518) | **706** | (25-3,000) | 0.1147 |
| **Total** | **11,245** | (3,643-22,505) | **15,605** | (8,204-42,156) | **16,131** | (5,653-55,345) | <.0001 |

In multivariate analysis, 4 factors were independently associated with higher direct medical costs: diagnosis on the basis of clinical signs, invasive cancer, lymph node involvement and conventional hospitalization for surgery (Table 4).

**Table 4:**
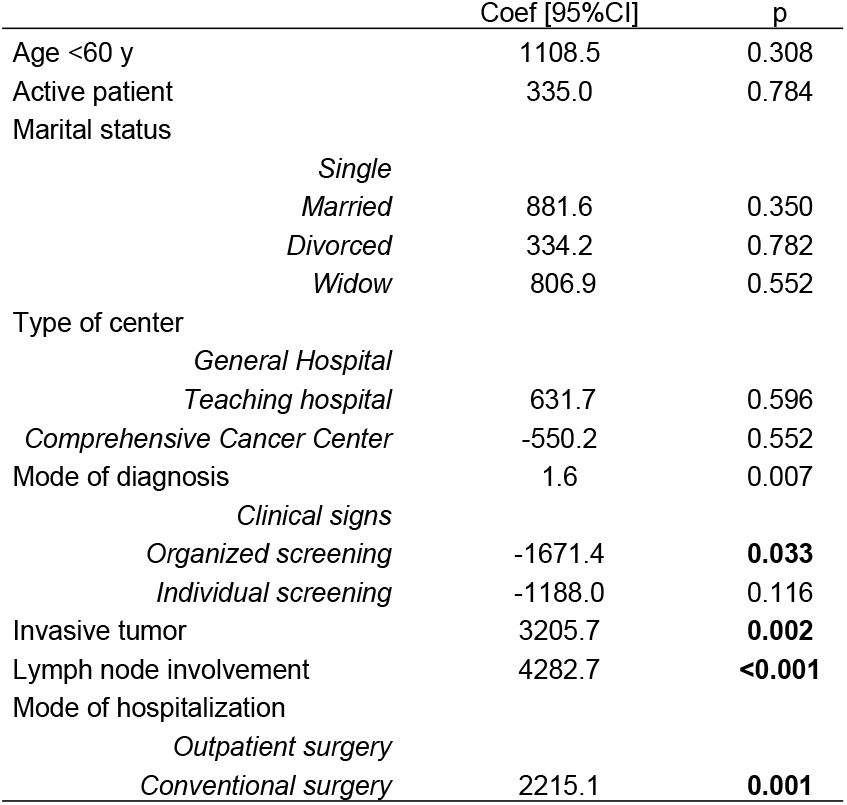
Multivariate analysis of the determinants of direct medical costs (multiple linear regression)

#### Direct non-medical costs from a National Health Insurance’s perspective

Sick leave costs were calculated for the 156 working patients included in the study for whom salary and duration of sick leave during the first year after diagnosis were available. Median sick leave costs were €8,841 per patient and per year (€173-€32,015). Only chemotherapy was found to be an independent determinant of sick leave costs, which were increased nearly threefold in the case of adjuvant chemotherapy (€3,687/patient/per year without chemotherapy versus €10,706/patient/year with chemotherapy, p<0.001).

One hundred and ninety-five patients declared transportation costs The median cost of transportation for these patients was €902 (€15 - €10,024). No significant difference in terms of transportation costs was observed between the 3 groups, and the only determinant identified on multivariate analysis was a distance between the patient’s residence and hospital greater than 16 km (OR=2.9; 95%CI[1,2-6,7]; p=0.01).

#### Direct medical and non-medical costs from a patient’s perspective

Forty percent (n=242) of patients declared additional personal expenditure, including medical (medical consultations and imaging) and non-medical costs (purchase of a wig, transportation, bras for prosthesis, etc.), with a median cost of €614 per patient and per year (€10-€16,909). Costs were higher for group 2 and 3 patients (€904 and €956, respectively) compared to group 1 patients (€363, p<0.001). In multivariate analysis, no additional factor was found to be associated with personal expenditure in this population.

## DISCUSSION

This multicenter prospective study describes BC pathways in a given health territory and the direct medical and non-medical costs during the first year after diagnosis. BC pathways mostly depend on 2 factors: initial stage of disease, which determines treatment and the resources consumed and the type of center. For example, more than 80% of group 1 patients undergoing breast-conserving surgery without adjuvant chemotherapy were operated on an outpatient basis. However, although this patient group is homogeneous, differences were observed in terms of health organization according to the type of center.

Undergoing surgery in a CCC was associated with outpatient surgery. Moreover, most patients of this study were managed in a CCC and we also observed that patients were willing to travel many miles to receive treatment in a CCC. All public centers involved in the management of BC in the area studied participated in this study; patient recruitment therefore reflects the reality of the health care territory. We have also previously shown that compliance with selected EUSOMA (European Society of Mastology) quality criteria was observed more frequently in CCC [17]. The type of center was therefore identified as an efficiency lever to optimize BC pathways. More BC patients are managed in CCC each year than in certain TH or GH, and they have a particular expertise in the management of cancer. Consequently, organization of care is optimized and can be easily be adapted when innovative technics or services become available, accounting for the high rate of outpatient BC surgery. The number of patients treated each year has been recently shown to be independent factor of overall survival in breast and ovarian cancer patients in a French National Health Insurance report comparing survival and the annual number of patients managed [18]. Women operated in a center performing less than 30 cases of BC surgery per year had an 84% excess risk of death compared to women operated in a center performing more than 150 cases of BC surgery per year.

Median direct medical costs of 1 year of management from diagnosis of operable BC were €12,250, from a National Health Insurance perspective. The cost of breast cancer in Europe has been estimated to be €15 billion [19]. In France, two recent studies evaluated the cost of breast cancer. The first study proposed a method based on formal concept analysis to form groups of hospital care trajectories [20]. By using an anonymous identifier to link sequences in different hospitals, the authors re-attributed all diagnostic codes to each patient admitted for surgery in 2009 and retraced their care pathway over a period of one year. The 57,552 patients selected over a 2-year period from data derived from the *Programme de médicalisation des systèmes d’information* (PMSI) [medical information systems programme] were classified into 19 care pathway groups, corresponding to particular morbidity profiles, as evaluated by main and associated diagnostic codes: invasive or *in situ* cancer, palliative care, aplasia, inflammatory breast disease. The average cost of a care pathway was €9,600. The greatest cost variability was explained by the invasive cancer code, associated with chemotherapy followed by palliative care, reconstructive surgery and aplastic anemia. The largest group in terms of numbers (62% of patients) had the lowest pathway cost. The second study proposed a macroeconomic method to calculate the overall cost of primary treatment of non-metastatic breast cancer using aggregated data from hospital activity, the national cost scale, and incidence data for a given year [21]. The average cost of the pathway without reconstituting individual pathways was €14,555 in 1999. These results were compared to a micro-economic method of cost evaluation, based on cost accounting. The pathway cost for 115 patients completely treated in a regional cancer center from diagnosis to the end of primary treatment was €14,399, a difference of only 1.1% compared to the cost estimated by the macroeconomic method. Similar results were recently obtained in Italy, based on cancer registry data [22]: mean total direct medical cost was €10,970/patient with €414 for diagnosis, €8,780 for treatment and €2,351 for 2-year follow-up, respectively.

These estimates are close to the direct medical costs estimated in the present study. However, these studies did not determine direct non-medical costs, such as transportation or sick leave costs. Nevertheless, median sick leave costs were €8,841/year/working patient, corresponding to the majority of absenteeism (National Health Insurance perspective) in the global cost of BC, and accounting for more than half of annual direct medical costs. In the light of these results, health authorities should develop individual support for working patients in order to decrease the duration of sick leave and improve return to work. Transportation costs, as well as out-of-pocket costs, were declared by a small proportion of patients and represented a minor share of global costs. No significant difference, especially in terms of sociodemographic characteristics, was observed between patients according to whether or not they reported out-of-pocket expenditures. Out-of-pocket expenditure obviously depends on the country of care. In France, cancer patients receive 100% cover of all breast cancer-related health care costs from French National Health Insurance. Out-of-pocket costs therefore essentially consist of complementary paramedical therapies (relaxation therapy, acupuncture, osteopathy), or additional prosthetic material (additional bras for external prosthesis, additional wig, etc.). Similar results were reported in Canada, with a median out-of-pocket expenditure of US$1002 during the year after diagnosis [23]. The burden on households depends on the household income. In the Optisoins01 study, nearly one-half of patients earned more than €1,900 per month [24]. An annual out-of-pocket expenditure of €614 therefore represents a reasonable burden for these households. Conversely, in Haiti, for example, the median non-medical out-of-pocket expenditure of BC patients was US$233 (from diagnosis to follow-up) for a median daily salary of US$2 [25]. In contrast, health care costs in some countries are not covered by national health insurance and patients need to take out private insurance. Out-of-pocket expenditure can therefore become a barrier to care. For example, in the US, the financial impact of BC has been described to prevent underserved BC survivors from accessing follow-up care [26].

A variability of direct costs is also observed between countries. Most studies in the US have described higher direct medical costs. A study based on a private insurance database reported a total health care cost of US$42,401 per patient for the first year [27]. In another study of patients covered by private insurance in the US, the annual direct medical cost of BC was US$19,435 [28]. Moreover, expenses as high as US$5,711 per month have also been published for patients covered by the Medicaid system [29]. However, in these last 2 studies, the study populations consisted of younger women aged 18-44 years. Age is a recurring determinant of costs [22,30], but was not significant in our study. The main determinant of cost in Optisoins01 was administration of chemotherapy, which accounts for the major difference in terms of direct medical costs between our BC pathway groups. A recent study also identified chemotherapy as one of the major drivers of direct medical costs, not only in BC, but also in lung and colorectal cancer [30].

BC costs mainly depend on treatment and BC pathways. Optimizing clinical pathways could therefore help to control costs. Moreover, implementation of organized clinical pathways helps to improve quality of care [31]. In the light of these results, our institute has established a quality charter involving several aspect of BC management and centered on clinical pathways to help other centers achieve excellence.

## Acknowledgments

This study was supported by a grant from the French National Cancer Institute (Institut National du Cancer, PRME-K2013), dedicated to economic studies of innovative techniques.

